# Exploitation of sulfated glycosaminoglycan status for precision medicine of platinums in triple-negative breast cancer

**DOI:** 10.1101/2021.05.05.442616

**Authors:** James D. Hampton, Erica J. Peterson, Samantha J. Katner, Tia H.Turner, Mohammad A. Alzubi, J. Chuck Harrell, Mikhail G. Dozmorov, Pam J. Gigliotti, Vita Kraskauskiene, Mayuri Shende, Michael O. Idowu, Madhavi Puchalapalli, Bin Hu, Larisa Litovchick, Eriko Katsuta, Kazuaki Takabe, Nicholas P. Farrell, Jennifer E. Koblinski

**Author notes:** These authors contributed equally to this study (James D. Hampton, Erica J. Peterson). Co-corresponding authors, these authors contributed equally as senior authors in the development of this study (Nicholas P. Farrell, Jennifer E. Koblinski). **Corresponding Author Information**: Jennifer Koblinski, PhD, Assistant Professor, Director, Mouse Model Core, Department of Pathology.

## Abstract

Triple negative breast cancer (TNBC) is a subtype of breast cancer lacking targetable biomarkers. TNBC is known to be most aggressive, and when metastatic is often drug resistant and uncurable. Biomarkers predicting response to therapy improve treatment decisions and allow personalized approaches for TNBC patients. This study explores sulfated glycosaminoglycan (sGAG) levels as a predictor of TNBC response to platinum therapy. sGAG levels were quantified in three distinct TNBC tumor models including cell line-derived, patient-derived xenograft (PDX) tumors, and isogenic models deficient in sGAG biosynthesis. The *in vivo* antitumor efficacy of Triplatin, a sGAG-directed platinum agent, was compared in these models to the clinical platinum agent, carboplatin. We determined that >40% of TNBC PDX tissue microarray samples have high levels of sGAGs. The *in vivo* accumulation of Triplatin in tumors as well as antitumor efficacy of Triplatin positively correlated with sGAG levels on tumor cells, whereas carboplatin followed the opposite trend. In carboplatin-resistant tumor models expressing high levels of sGAGs, Triplatin decreased primary tumor growth, reduced lung metastases and inhibited metastatic growth in lungs, liver, and ovaries. sGAG levels served as a predictor of Triplatin sensitivity in TNBC. Triplatin may be particularly beneficial in treating patients with chemotherapy-resistant tumors who have evidence of residual disease after standard neoadjuvant chemotherapy. More effective neoadjuvant and adjuvant treatment will likely improve clinical outcome of TNBC.

**Significance:** TNBC is a heterogenous disease, often defined by the absence of a therapeutic target. Our recent results show sGAGs may provide a viable biomarker for Triplatin in patients with TNBC, producing a significant advantage over carboplatin in this setting. Selective precision medicine agents, such as Triplatin, that are active against chemotherapy-resistant disease and exploit molecular biomarkers like sGAGs may significantly benefit patients in this setting.

## Introduction

Triple negative breast cancer (TNBC) represents 15-20% of breast cancers and is defined by the lack of estrogen-and progesterone-receptor expression and absence of overexpression of human epidermal growth factor receptor 2 (HER2). Chemotherapy remains the primary treatment, and the Standard of Care is to use taxanes and anthracyclines as first-line therapy for both early stage and metastatic TNBCs (1,2). TNBC is known to be the most aggressive subtype and recurs in half of patients after curative treatment, and rapidly develops drug resistance and metastases. The average life expectancy of approximately 1 year after recurrence underscores the need for novel therapeutic regimens that can improve survival outcomes (3).

Recent clinical trials investigating strategies to optimize neoadjuvant and adjuvant treatment options include use of immunotherapy, platinum agents and PARP inhibitors (4). Trial results for early stage TNBC show that addition of carboplatin or carboplatin/pembrolizumab to standard neoadjuvant chemotherapy (NACT) regimens significantly increases the number of patients who achieve a pathological complete response (pCR), which is a known surrogate for better survival (4,5). For metastatic TNBC expressing PD-L1, the FDA recently approved a combination of nab-paclitaxel and atezolizumab, which showed improved progression-free survival (PFS) in the IMpassion130 trial (6,7). Furthermore, the Phase III TNT trial showed higher overall response rates (68% vs 33%; p=0.01) and a longer median PFS in patients with metastatic TNBC harboring BRCA mutations who received carboplatin presented in comparison with those who received docetaxel (8). However, the need for efficient treatments remains high and biomarkers predicting response to therapy could help to inform treatment decisions and allow for personalized approaches for TNBC patients.

Studies aimed at the identification of biomarkers and treatment targets of TNBC have mainly focused on mRNAs (9), miRNAs, and proteins, while carbohydrates, such as glycosaminoglycans (GAGs), are relatively unexplored. The non-template driven nature of GAG biosynthesis involves the concerted action of many enzymes, which complicates the analysis of expression at the genomic/proteomic level and their role in clinical oncology. GAGs, such as heparan sulfate (HS) and chondroitin sulfate (CS) are linear polymers of repeated disaccharide units that are modified with sulfate groups at various positions. The resulting sulfated GAGs (sGAGs) are highly negatively-charged and are known to mediate interactions of angiogenic factors, such as cytokines and growth factors with their respective cell surface receptors (10). Changes in chain length, sulfation, and the expression of proteoglycans (containing GAG attachment sites) at the cell surface and within the extracellular matrix are associated with the progression of TNBC and other cancers (11,12) (Table 1).

**Table 1.** Overexpression of sulfated glycosaminoglycans in breast cancer

| Observation | Method | Sample type | Ref |
| --- | --- | --- | --- |
| ↑ in total sulfated GAGs | carbazole, PAGE | Clinical (BC) | (23) |
| ↑ in total sulfated GAGs | Alcian Blue (IHC), DMMB | Clinical (BC), MDA-MB-231 | (24) |
| ↑ N-sulfated HS | Ab 10E4 (IHC) | Clinical (BC, TNBC) | (25-27) |
| ↑ N-sulfated HS | Ab 10E4 (flow cytometry) | MDA-MB-231, HCC38 | (28) |
| ↑ sulfated CS | Ab CS-56 (IHC, flow cytometry) | Clinical (BC), MDA-MB-231 | (29) |
| ↑ sulfated CS | VAR2CSA | Clinical (BC), MDA-MB-231 | (30) |

Recently, we discovered that the positively-charged platinum agent, Triplatin (BBR3464), has high affinity for sGAGs, which mediates its cellular accumulation and cytotoxicity, while these properties are not shared by either cisplatin or carboplatin (13). Triplatin-sGAG interactions inhibit cancer cell growth and invasion by blocking growth factor signaling and heparanase activity (14). In this contribution, we explore sGAGs as a predictive marker for Triplatin efficacy against models of TNBC primary tumor growth and metastasis.

## Materials and Methods

### Reagents

Triplatin was synthesized by published procedures (15). Carboplatin and cisplatin were purchased from Sigma.

### Cell lines and culture conditions

CHO-K1, CHO pgsE-606, and CHO pgsA-745 were purchased from ATCC. CHO pgsF-17 was provided by Dr. Jeffrey Esko at the University of California San Diego. All CHO cell lines were cultured in DMEM-F12 media with 10% fetal bovine serum (FBS) and 1% penicillin/streptomycin. MDA-MB-231-BrM2-*luc* cells (provided by Dr. Joan Massauge at Sloane Kettering Cancer Center) and MDA-MB-231-*luc* cells (provided by Dr. Danny Welch at University of Kansas) were cultured in DMEM media (Invitrogen) with 10% FBS, 1% L-glutamine, and 1% penicillin/streptomycin (Gibco). The MDA-MB-231-luc cells were generated by infection with lentiviruses containing the pFULT vector expressing fire-fly luciferase. The lentiviruses were generated by DNA/RNA Delivery Core, Skin Disease Research Center, Department of Dermatology, Northwestern University Feinberg Medical School. 4T1-*luc* cells (provided by Dr. Sarah Spiegel at Virginia Commonwealth University) were cultured in RPMI media with 10% FBS and 1% penicillin/streptomycin. All cells were grown in a humidified atmosphere at 37°C with 5% CO_2_. Cells were routinely checked for mycoplasma contamination using the mycoplasma detection kit (Lorenzo).

### Generation of Xylt1/2 knock-out cell lines

sGAG deficient MDA-MB-231 cell lines were generated by transfecting the parental/wild-type MDA-MB-231-luc cell line according to the protocol found in (16). Cells received either Cas9 plasmid and empty DNA backbone (control cells) or Cas9, *Xylt1*, and *Xylt2* gRNA plasmids (Knock-out [KO] cells). Following transfection, cells were single cell sorted based upon GFP expression and allowed to grow in 96-well plates for 4 weeks, to allow for the clonal expansion of individual cell lines. Wild-type (wt) and generated control, and KO cells were then screened *via* flow cytometry using sGAG specific antibodies, to detect sGAG deficient cell lines with XYLT1/2 KO and to confirm that control cell lines matched wild-type sGAG levels. Following screening, cells were used for *in vitro* and *in vivo* studies. Genomic alterations in KO cells were confirmed via cloning of Xylt1/2 PCR products and ligation into TOPO TA sequencing plasmid. Sequencing was carried out using Euro Fins Sanger Sequencing, M13F primer. PCR Primers – Xylt1fwd (gtttctctcccctcttctccag), Xylt1rev (atgtcacacttagggggctg), Xylt2fwd (cccagtgatgtgtctgcatc), Xylt2rev (gtatctccgtgtggtgggaag). Each cell line was sequenced 6 times for each gene to ensure accuracy.

### sGAG quantitation

*Blyscan assay*. The Blyscan assay was used per the manufacturer’s instructions (Biocolor). Cells were seeded onto 100 mm dishes and grown to 90% confluence. Cells and cellular matrix were dissociated from dishes in 1 ml trypsin-EDTA and placed directly on ice. After the number of cells in each sample were determined, 100 μl of the suspension was added to 1 ml of 1,9-dimethylmethylene blue (DMMB) dye reagent and mixed *via* inversion every 5 min for 30 min. The samples were centrifuged at 12,000 rpm for 10 min. to pellet precipitated GAG-dye complexes. Complexes were then resolved in SDS-containing dissociation reagent and the absorbance of bound dye was measured at 656 nm. Values were compared to chondroitin-4-sulfate standards and normalized to cell number.

*Flow cytometry*. To study the relative amounts of cell surface sGAGs expressed, the binding of the 10E4 antibody (clone F58-10E4, Amsbio) to sulfated HS can be quantitated using flow cytometry. Briefly, cells are harvested using versene and washed twice in cold PBS. 1.5 × 10^6^ cells are resuspended in PBS containing 0.5% BSA and kept on ice throughout the entire procedure. The cells are incubated with 1:100 10E4 primary antibody for 30 min. After washing twice with PBS, the cells are incubated with 1:400 goat anti-mouse IgM Alexa-647 conjugated secondary antibody (Life Technologies). Cells incubated with only secondary antibodies serve as negative controls. Fluorescence is measured using a Fortessa flow cytometer (BD Biosciences).

*Immunohistochemistry (IHC)*. All tissues were fixed in 10% neutral buffered formalin for at least 5 days before paraffin embedding and sectioning. A tissue microarray (TMA) was obtained from Champions Oncology containing breast cancer PDX samples. IHC staining for human leukocyte antigen (HLA) primary antibody (Abcam, clone EMR8-5, 1:50 dilution) and N-sulfated heparan sulfate (Amsbio, clone F58-10E4, 1:100 dilution) was performed in the VCU Cancer Mouse Models Core with the Leica Bond RX autostainer. All antibodies were incubated for 30 min at 25°C. For HLA staining heat-induced epitope retrieval buffer 2 (Leica, EDTA pH 8.0) was used. For negative control of N-sulfated heparan sulfate staining (10E4), slides were incubated with 50 mU Heparinase III (Sigma-aldrich) in Tris buffer (20 mM Tris-HCL, pH 7.5 containing 0.1 mg/ml bovine serum albumin and 4 mM CaCl2) for 45 min at 37°C, 3 times, before primary antibody incubation. Digestion with heparinase III cleaves the heparan sulfate chain and results in a loss of the 10E4 epitope. For the positive N-sulfated heparan sulfate staining (10E4) slides were incubation with the Tris buffer only for 45 min at 37°C, 3 times, before primary antibody incubation. Bond polymer refine detection kit was used for DAB chromogen and hematoxylin counterstain (Leica). The tissue slides were visualized with a Vectra^®^ Polaris™ Automated Quantitative Pathology Imaging System (Akoya Biosciences) using whole-slide scanning at 20x. Phenochart Whole Slide Contextual Viewer software was used to visualize the scans and inFrom software (Akoya Biosciences) was used to quantify the staining (17–19). The CHOK1 and psA-745 were used to titrate the 10E4 antibody and validate staining. Liver metastases were quantified by metastases cell number identified by the HLA staining using Cell Counter on ImageJ and then normalized to total liver area measured in ImageJ. n = 5, each group.

### Cell Growth Inhibition

One thousand cells/well were seeded in 96-well plates. After 24 h, cells were treated with either carboplatin or Triplatin for 15 min in 200 μl media. Following treatment, drug media was removed, wells were washed with phosphate buffered saline (PBS) and replaced with drug-free media. Images and confluence data were acquired every 2 h for ~200 h using the IncuCyte Live Cell Analysis Imaging System (Sartorius). The Area Under Curve (AUC) was calculated for each concentration using GraphPad Prism 8.0, normalized against the control, and used to calculate the IC50 values for Triplatin and carboplatin.

### Cellular Platinum Uptake

Intracellular platinum content was determined as previously described (13). Cells were seeded onto 100 mm dishes, allowed to attach for 24 h, and treated with 10 μM carboplatin or Triplatin. Following 1 or 3 h incubations, cells were harvested *via* trypsinization and counted. Cell pellets were washed repeatedly with PBS and digested in nitric acid for 72 h. Samples were filtered and analysed by inductively-coupled plasma mass spectrometry (ICP-MS). Values were compared to platinum standards and normalized to cell number.

### Tumor Platinum Uptake

CHO-K1 and pgsA-745 (3.0 × 10^6^) cells were suspended 1:2 with 100 µl of PBS:14 mg/ml of Cultrex BME Type 3 (Bio-Techne) and subcutaneously injected into the left and right flanks of male NSG mice, respectively as described in (20). MDA-MB231 Xylt1/2 KD cells (1.0 × 10^4^) were injected intraductally into female NSG mice. Once the tumors grew to a size of ~400 mm^3^, mice were given a single dose of Triplatin (0.3 mg/kg) or carboplatin (40 mg/kg) and euthanized 24h after administration. Tumors were harvested, weighed, dissolved in nitric acid for 72 h, and filtered. Platinum content was determined using ICP-MS as previously described (13). Values were normalized to wet tumor weight.

### Drug Efficacy

*In vivo mouse models*. Virginia Commonwealth University Institutional Animal Care and Use Committee (IACUC) approval was obtained for all experiments. NOD.Cg.-Prkdc^scid^IL2rg^tm1Wjl^/SzJ (NSG) mice were purchased from the Cancer Mouse Models Core Laboratory at VCU Massey Cancer Center. Balb/c mice were purchased from Envigo.

*Orthotopic primary tumor studies*. MDA-MB-231-luc cells (1×10^4^), WHIM2-luc, UCD52-luc, and WHIM30-luc (1×10^5^) PDX cells were suspended in 100 µl of a 1:9 mix of PBS:Cultrex BME Type 3 (14 mg/ml, Bio-Techne) and injected into the 4^th^ lactiferous duct of 6-8-week-old female NSG mice as described in (21,22). For 4T1 studies, (1×10^4^ cells suspended in 20µL 1:9 PBS: Cultrex BME Type 3) were injected in the 2^nd^ lactiferous duct of 6-8 week old Balb/c mice. Drug treatments (vehicle [saline], cisplatin [3 mg/kg], carboplatin [40 mg/kg], or Triplatin [0.3mg/kg]) were given interperitoneally (i.p.) on days 1, 5, and 9 (Q4Dx3) after randomization with day 1 being the day of randomization. Tumor growth was measured by using either digital caliper or bioluminescent imaging with the IVIS Spectrum (PerkinElmer). Using Studylog lab management software, the mice were randomized with the 2 factor (tumor volume and body weight) randomization Multi-Task program into groups once tumors reached an average of 150 mm^3^. At the endpoint, all mice were imaged live, euthanized and tumors were harvested and weighed. The bioluminescence of harvested tumors and organs were quantified by *ex vivo* imaging and analysis. Each organ was immersed for 5 min in luciferin/PBS (0.3 mg/ml).

### Experimental metastasis studies

MDA-MB-231-BrM2-*luc* cells (1×10^5^ for endpoint study or 1×10^4^ for survival study) or WHIM2-luc, UCD52-luc, and WHIM30-luc (5×10^5^ for endpoint studies) PDX cells in 100 µl sterile PBS were injected into the left cardiac ventricle of female 5-week-old NSG mice as described (22). The cells were mixed with luciferin (1.5 mg/ml) to permit immediate *in vivo* imaging (IVIS Spectrum, Perkin Elmer) to verify widespread seeding of tumor cells. Mice were randomized using Studylog lab management software with the 2 factor (total flux values and body weight) randomization Multi-Task program on day 10. The groups were as follows: 2 groups (control and Triplatin) for MDA-MB-231-BrM2-*luc* cells endpoint studies or 3 groups (vehicle, carboplatin, Triplatin) for WHIM2-luc, UCD52-luc, and WHIM30-luc endpoint studies, or 3 groups (vehicle, cisplatin, and Triplatin) for MDA-MB-231-BrM2-*luc* cells survival studies. After randomization mice were treated i.p. with saline vehicle, cisplatin (3 mg/kg), carboplatin (40 mg/kg) or Triplatin (0.3 mg/kg) on days 1, 5 and 9 after randomization with day 1 being the day of randomization. Tumor burden was quantified by bioluminescence (radiance/sec) emitted from the cells after a subcutaneous injection of luciferin (150 mg/kg diluted in PBS) allowing for live *in vivo* imaging using the IVIS Spectrum and Living Image® software (PerkinElmer). For the endpoint study, mice were euthanized and organs (kidneys, heart, lung, ovaries, liver, brain, and skeleton) were harvested for *ex vivo* imaging as described above.

### Statistical analysis

One-way analysis of variances (ANOVA) was performed to determine statistically significant differences among the control and treatment groups. Tukey’s multiple comparisons tests were performed to compare across all groups. Unpaired t-tests were performed to compare 2 groups. Kaplan-Meier survival plots were generated and changes in survival were analyzed by the log-rank test. All analyses were calculated in GraphPad Prism 8 software and p values < 0.05 were considered significant.

## Results

### Sulfated glycosaminoglycan profile may be exploited for therapeutic benefit in TNBC

Dysregulation of sGAG expression affects all stages of tumorigenesis and correlates with poor prognosis in multiple cancers (11,12). These findings have raised considerable interest in the development of sGAG-based qualitative and quantitative methods which include: alcian blue, 1,9-dimethylene blue (Blyscan), capillary electrophoresis, high-performance liquid chromatography (HPLC), mass spectrometry, and IHC. These methods have all been used to determine the relevance and significance that sGAGs play in biological processes, as identification of changes at the RNA/protein level are not sufficient to accurately predict changes in cell sGAG profile. Several reports observed increases in sGAG levels in TNBC and other BC cell lines as well as clinical samples compared to normal tissue, Table 1.

Based on these observations, we analyzed a TMA of TNBC PDX samples for sGAG levels using two GAG-specific antibodies; Ab 3G10, which reacts with an HS neoepitope after heparinase III digestion and Ab 10E4, which reacts with N-sulfated HS. The Ab 10E4 staining showed a wide range of expression, with two distinct populations staining high (40%) and low (60%) (Fig. 1A,1B), whereas Ab 3G10 staining was high overall and less dynamic (Fig. S1). These results confirmed that sGAGs are upregulated in TNBC and thus, sGAG levels may be exploited for precision medicine with a compatible therapeutic.

**Figure 1.**
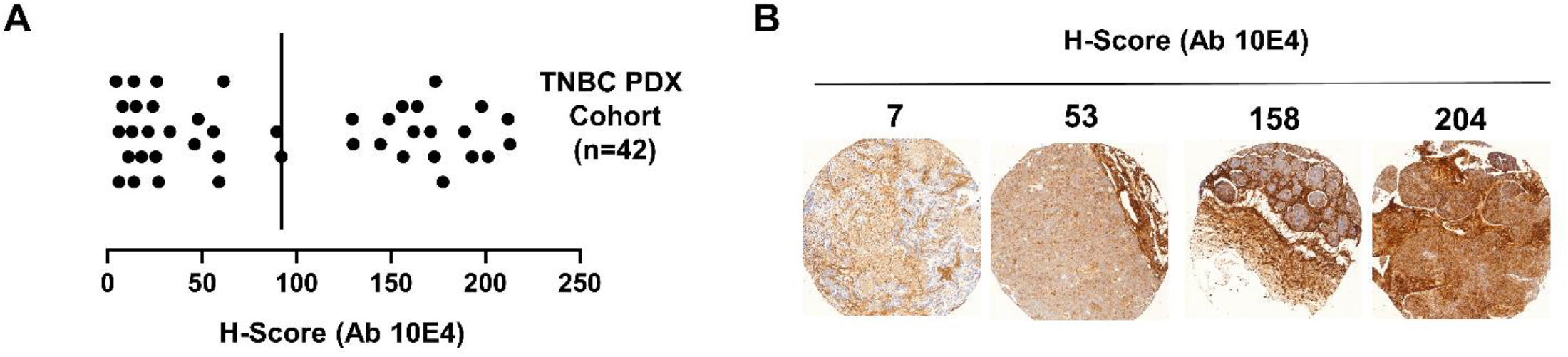
Sulfated glycosaminoglycans are a potential target for Triplatin in TNBC. **A**. IHC H-scores of TNBC PDX samples on TMA (n= 42) stained with Ab 10E4. Each data point represents the mean H-score for duplicate samples. **B**. Representative images of Ab 10E4 staining of TNBC PDX TMA samples.

### Triplatin tumor uptake is sGAG-dependent

The positively-charged polynuclear platinum agent, Triplatin (Fig. 2A) interacts with negatively-charged sGAGs with high affinity. The optimized structure for the interaction of [Pt(NH3)4]2+, the central platinum of Triplatin, with a sulfated monosaccharide, GlcNS(6S) is shown in Fig. 2B (31). Wild-type (wt) CHO-K1 cells and GAG-deficient CHO mutant cell lines pgsF-17, pgsE-606, and pgsA-745 (Fig. 2C) were used to determine how sGAG levels affect platinum uptake. The pgsF-17 (F17) cell line, harboring a mutation in HS 2-0-sulfotransferase (HS2ST1), does not show significant changes in total sGAG levels as measured by DMMB-based Blyscan assay (Fig. 2D), and instead shows a slight compensatory increase in N-sulfated HS (Fig. 2E). The pgsE-606 (E606) cell line, with a mutation in HS N-sulfotransferase (NDST1), and the pgsA-745 (A745) cell line, with a mutation in one of the rate-limiting controller of sGAG biosynthesis, xylosyltransferase-2 (XYLT2), show a significant reduction in both total sGAG and N-sulfated HS (Fig. 2C-E, 2I). To determine the effect of sGAG levels on platinum uptake, we compared the cellular accumulation of Triplatin and carboplatin in these cell lines (Fig. 2F). The overall cellular accumulation of Triplatin in the control cell line was significantly higher than that of carboplatin, as expected from previous results (13). The cellular accumulation (Fig. 2F) and cytotoxicity (Fig. 2G-H) of Triplatin increases with increasing sGAG levels (IC50 decreases from 120 ± 4.2 nM to 69.5 ± 0.3 nM). Surprisingly, the cellular accumulation (Fig.2F) and cytotoxicity (Fig. 2G-H) of carboplatin negatively correlates with sGAG levels (IC50 increasing from 434 ± 71 nm to ≥ 1000 nM). Furthermore, significantly more Triplatin accumulated *in vivo* in the wt compared to the GAG-deficient A745 tumor, whereas carboplatin showed a slightly increased accumulation in the A745 tumor compared to the wt (Fig. 2I, K). These results raise the intriguing possibility that sGAGs not only mediate Triplatin uptake but also serve as a predictive biomarker for patient treatment.

**Figure 2.**
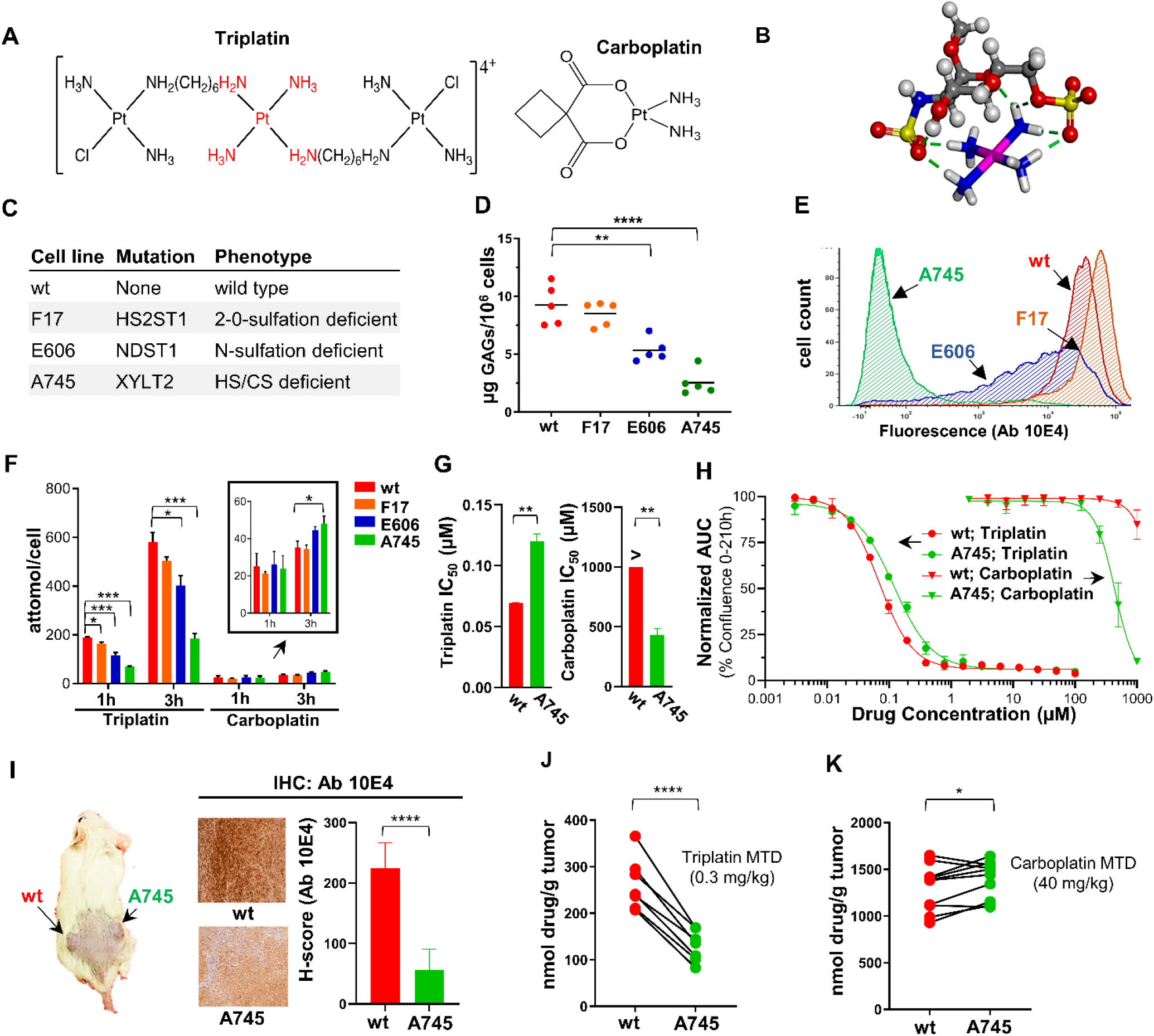
sGAG expression levels affect cellular accumulation, cytotoxicity, and tumor uptake of Triplatin. **A**. Structures of Triplatin and carboplatin **B**. DFT optimized structure for the interaction of [Pt(NH_3_)_4_]^2+^, the central platinum of Triplatin, with sulfated monosaccharide, GlcNS(6S). **C**. Mutation and phenotype of GAG-and sulfation-deficient CHO carcinoma cell lines **D**. Quantification of the total cell surface sGAG levels in CHO cell lines using the Blyscan sGAG quantitation assay (n=5 biological replicates). **E**. Relative levels of N-sulfated HS (Ab 10E4) expression by flow cytometry. Histogram representative of 3 independent experiments. **F**. ICP-MS quantification of Triplatin and carboplatin cellular uptake. **G-H**. Analysis of cell proliferation after 15 min treatments with the indicated concentrations of Triplatin or carboplatin. % confluence was quantified every 2 h for 210 h using live cell analysis (Incucyte). Each data point represents the mean ± SD (n=2 biological replicates with 4 technical replicates each). IC_50_ calculations are shown in G. **I**. Right panel, Image of a mouse injected with wtCHO and A475. Middle panel, IHC image of N-sulfated HS (Ab 10E4) expression in CHO wt and 745 tumors and staining was quantified (left panel). Each bar represents the mean ± SD (n=3). **J**,**K**. ICP-MS analysis of Pt tumor accumulation in CHO wt and A-745 tumors 24h after treatment with Triplatin (0.3 mg/kg; n=7) and carboplatin (40 mg/kg; n=10). Statistical analysis: D-F, One-way ANOVA, Tukey’s posttest; G unpaired student’s t-test; J and K, paired student’s t-test. *p<0.05, **p<0.01, ***p<0.001, ****p<0.0001.

### Triplatin has reduced accumulation in sGAG deficient TNBC cells and inhibits primary tumor growth and metastasis in carboplatin resistant TNBC models

To confirm these results in a more clinically relevant model, we generated XYLT1/2 knockout MDA-MB-231 cell lines using well characterized CRISPR plasmids as described (16). Created CRISPR clones KO-F and KO-G both possess Indels in Exon 3 of XYLT1 and Exon 2 of XYLT2, as determined by sanger sequencing (Fig. S3 A-B). These generated cell lines, MDA-MB-231 XYLT1/2 KO-F and KO-G, had >90% reduction in sGAG epitopes in comparison to both parental and control cells (Fig. 3A). Consistent with our model that high sGAG levels facilitate Triplatin accumulation, the MDA-MB-231 XYLT1/2 KO-F and KO-G cell lines had reduced uptake of Triplatin and showed higher uptake of carboplatin *in vitro* (Fig. 3B) compared to wt MDA-MB-231. Differences in Triplatin but not carboplatin accumulation were also seen *in vivo* (Fig. 3C-D). These results were similar to Triplatin and carboplatin accumulation in wt CHO K1 compared to CHO-pgsA-745 cells.

**Figure 3.**
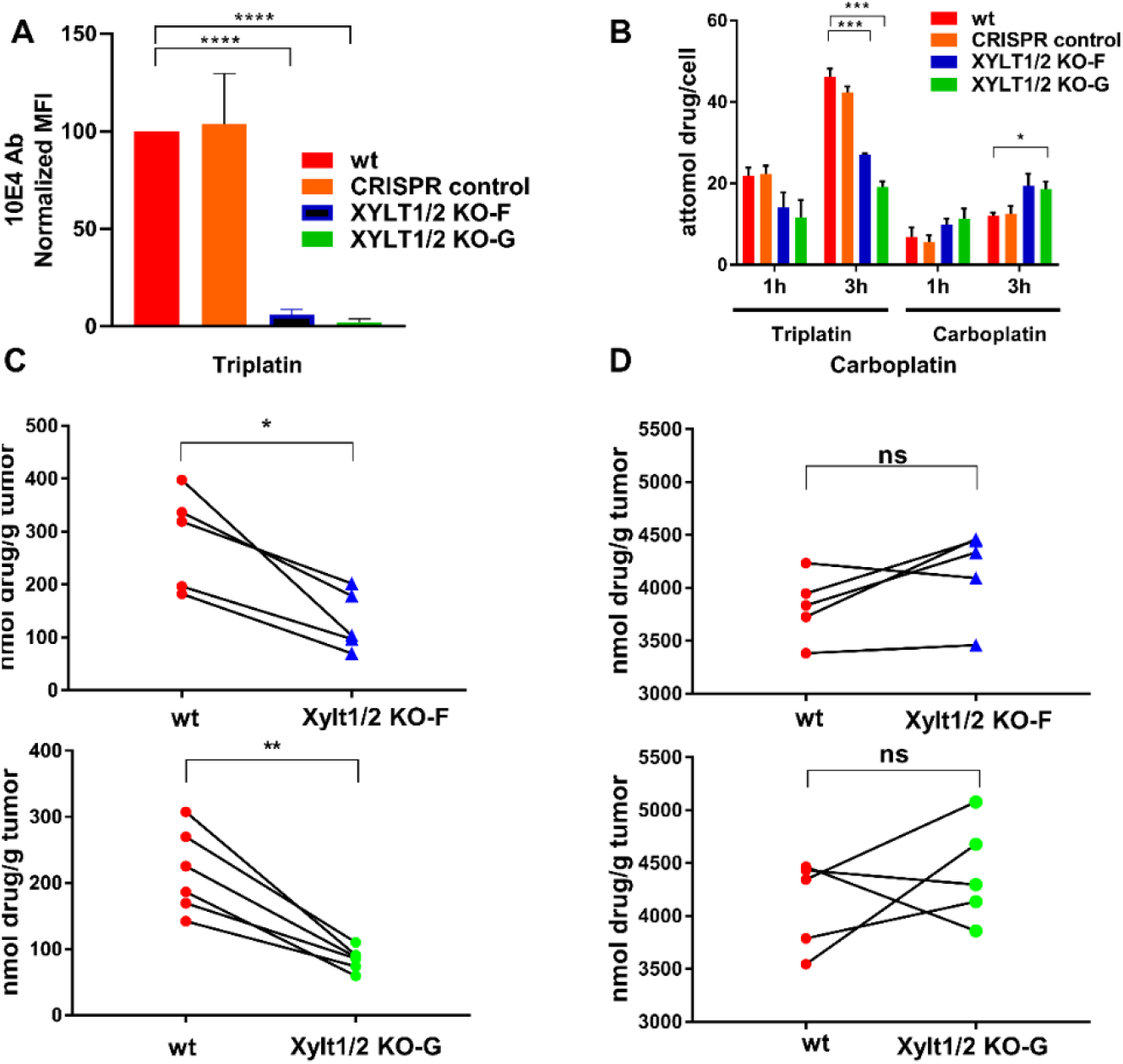
Cellular accumulation of Triplatin in MDA-MB-231 tumors is affected by sGAG levels. **A**. Flow cytometry analysis shows levels of N-sulfated HS (Ab 10E4) in MDA-MB-231 XYLT1/2 KO cell lines is significantly decreased compared to wt and CRIPSR control MDA-MB-231 cells. **B**. ICP-MS quantification of Triplatin and carboplatin cellular uptake *in vitro*. **C**,**D**. ICP-MS analysis of Pt tumor accumulation in parental MDA-MB-231 tumors and MDA-MB-231 XYLT1/2 KO-F and KO-G tumors, 24h after treatment with Triplatin (0.3 mg/kg; n=11) and carboplatin (40 mg/kg; n=10). Statistical analysis: A,B, One-way ANOVA, Tukey’s posttest. C-D, paired student’s t-test. *p<0.05, **p<0.01, ***p<0.001, ****p<0.0001.

Next, the effectiveness of platinum agents was compared against primary tumor growth using the parental MDA-MB-231-luc mouse TNBC model. MDA-MB-231 tumors show high reactivity with the 10E4 antibody (H-score >200), (Fig. 4A, B), comparable to that of the wild-type CHO tumors. To confirm the specificity of Ab 10E4 staining, the MDA-MB-231-luc tumor samples were incubated with heparinase III, which resulted in cleavage of the HS polymer and loss of the epitope against N-sulfated HS (Fig. 4A). To validate our hypothesis that Triplatin will be efficacious in tumor models with high levels of sGAGs, MDA-MB-231-luc cells were injected into the mammary gland of female NSG mice. Once tumors reached ~150 mm^3^, mice were randomized and treated with either saline (vehicle), Triplatin or carboplatin on days 1, 5, and 9 with day 1 being the day of randomization. As predicted, Triplatin was more effective than carboplatin at reducing tumor volume, tumor weight, and lung metastases in this model (Fig. 4C, D and S3A-B).

**Figure 4.**
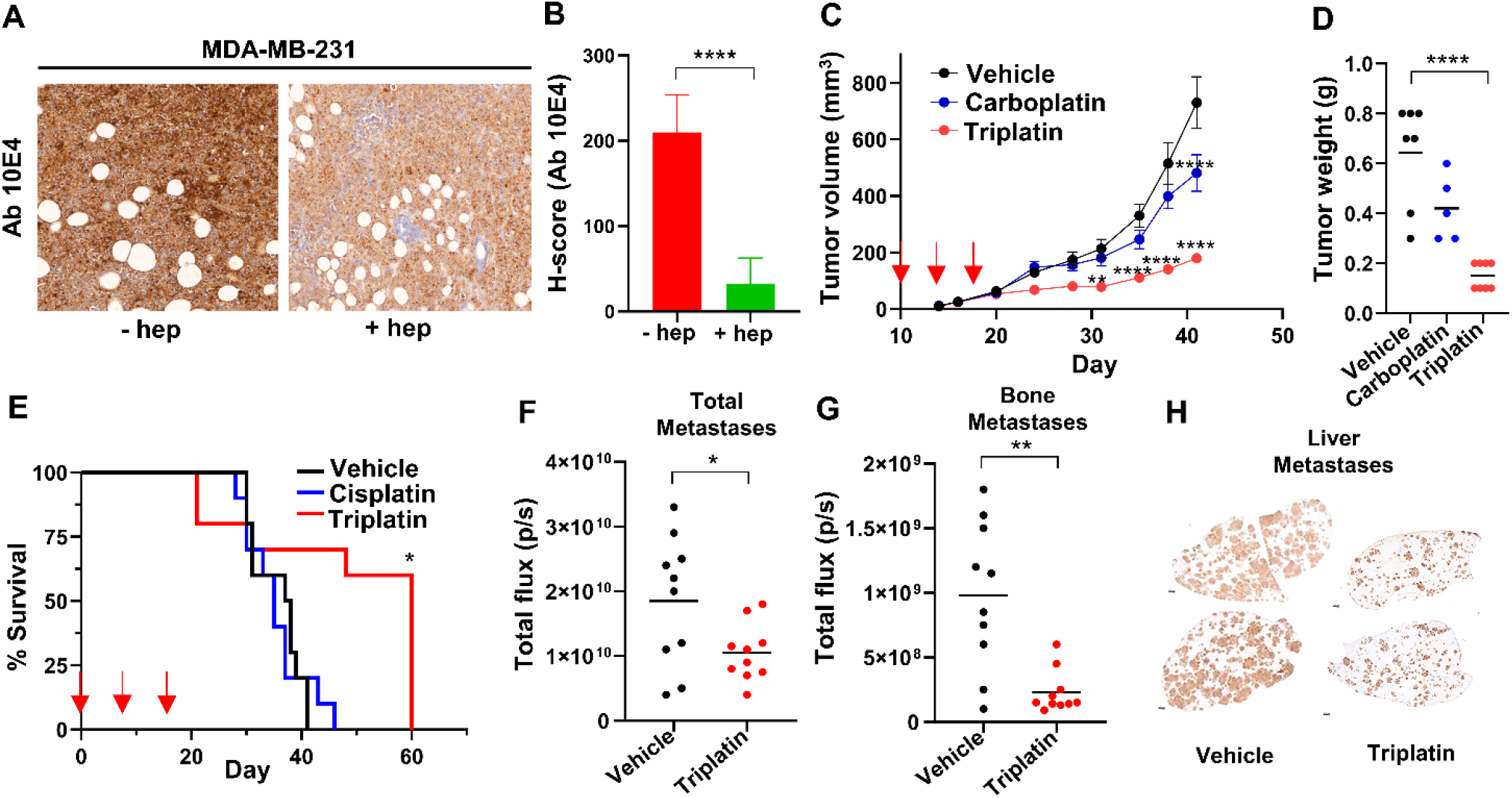
Triplatin inhibits tumor and metastatic growth in a high sGAG-expressing MDA-MB-231-luc model of TNBC. **A**. Representative images of IHC analysis of N-sulfated HS (Ab 10E4) reactivity in MDA-MB-231-luc tumors with or without heparinase III digestion. **B**. Quantitation of Ab 10E4 IHC (n=3). **C**. Analysis of primary tumor growth. Arrows, drug treatment [Triplatin (0.3mg/kg), carboplatin (40 mg/kg), or vehicle (saline)] given i.p. on days 1, 5, and 9 after randomization (day 16). **D**. Tumor weight at endpoint. **E**. Survival analysis: MDA-MB-231BR cells were injected into left ventricle of female NSG mice (day 0). Animals were randomized on day 10. Drug treatment [Triplatin (0.3mg/kg), cisplatin (3 mg/kg), or vehicle (saline)] given i.p. on days 10, 14, and 18. **F**. Quantification of overall metastases *in vivo* on day 21 (total flux). **G**. Quantification of bone metastases *ex vivo* on day 21 (total flux). **H**. Quantification of liver metastases *ex vivo* on day 21 using HLA staining. Statistical analysis: B, F and G, student’s t-test; C, D and E, One-way ANOVA; Tukey’s posttest *p<0.05, **p<0.01, ***p<0.001, ****p<0.0001.

Since the lower tumor burden can also result in decreased metastasis, we next determined the effect of Triplatin on metastatic growth using an intracardiac MDA-MB-231BR model of TNBC metastasis which allows analysis of Triplatin on inhibition of growth at metastatic sites seen in TNBC patients, e.g. the brain, bone, lung, and liver. Triplatin-treated mice gave marked increases in survival compared to control and cisplatin-treated mice (Fig. 4E). In an endpoint study, when all mice were sacrificed 21 days after cancer cell injection, total metastases and, especially, bone and liver metastases were dramatically reduced in mice treated with Triplatin compared to control mice (Fig. 4F-H, S4).

The ability of Triplatin to directly inhibit lung metastases was evaluated using the syngeneic murine 4T1 mastectomy model (32). The 4T1-luc mouse-derived mammary cancer cell line injected into immune-intact mice mimics human cancer progression and metastasizes more efficiently than conventional human xenografts. The IHC analysis of 4T1 tumors grown orthotopically showed high Ab 10E4 staining (Fig. S5A), suggesting that this model will be sensitive to Triplatin. Indeed, we observed that Triplatin inhibited lung metastases that occurred after mastectomy (Fig. S5B-C) and significantly increased survival (Fig. S5D). Furthermore, 4T1 cells were intracardially injected into mice to mimic widespread metastasis. Mice treated with Triplatin had reduced metastases in the lungs and bone compared with carboplatin-treated mice and saline control (Fig. S6A,B). Together, these results show that Triplatin inhibits primary tumor growth and metastasis in carboplatin-resistant TNBC models expressing high levels of sGAGs.

### The efficacy of Triplatin against TNBC PDX models correlates with sGAG levels

Use of PDX models in preclinical studies correlates well with clinical response (33–35). Therefore, the efficacy of Triplatin was compared in TNBC PDX with high and low levels of sGAGs to test the hypothesis that PDX models with high levels of sGAG staining will be most responsive to Triplatin. We examined 3 PDX TNBC models, previously investigated for sensitivity to carboplatin – UCD52 (carboplatin-sensitive), WHIM30 (carboplatin-sensitive) and WHIM2 (carboplatin-resistant) (36–39). WHIM30 and WHIM2 had significantly higher sGAG levels compared to UCD52 (Fig 5A-B). WHIM30 primary tumor growth was significantly inhibited by Triplatin and carboplatin compared to the control (Fig. 5C). Triplatin was more effective than carboplatin at inhibiting WHIM2 tumor growth (Fig. 5D), while carboplatin was more effective than Triplatin at inhibiting UCD52 primary tumor growth (Fig. 5E). These results indicate that the activity of Triplatin against primary tumor growth is correlated with sGAG levels in TNBC PDX tumors.

**Figure 5.**
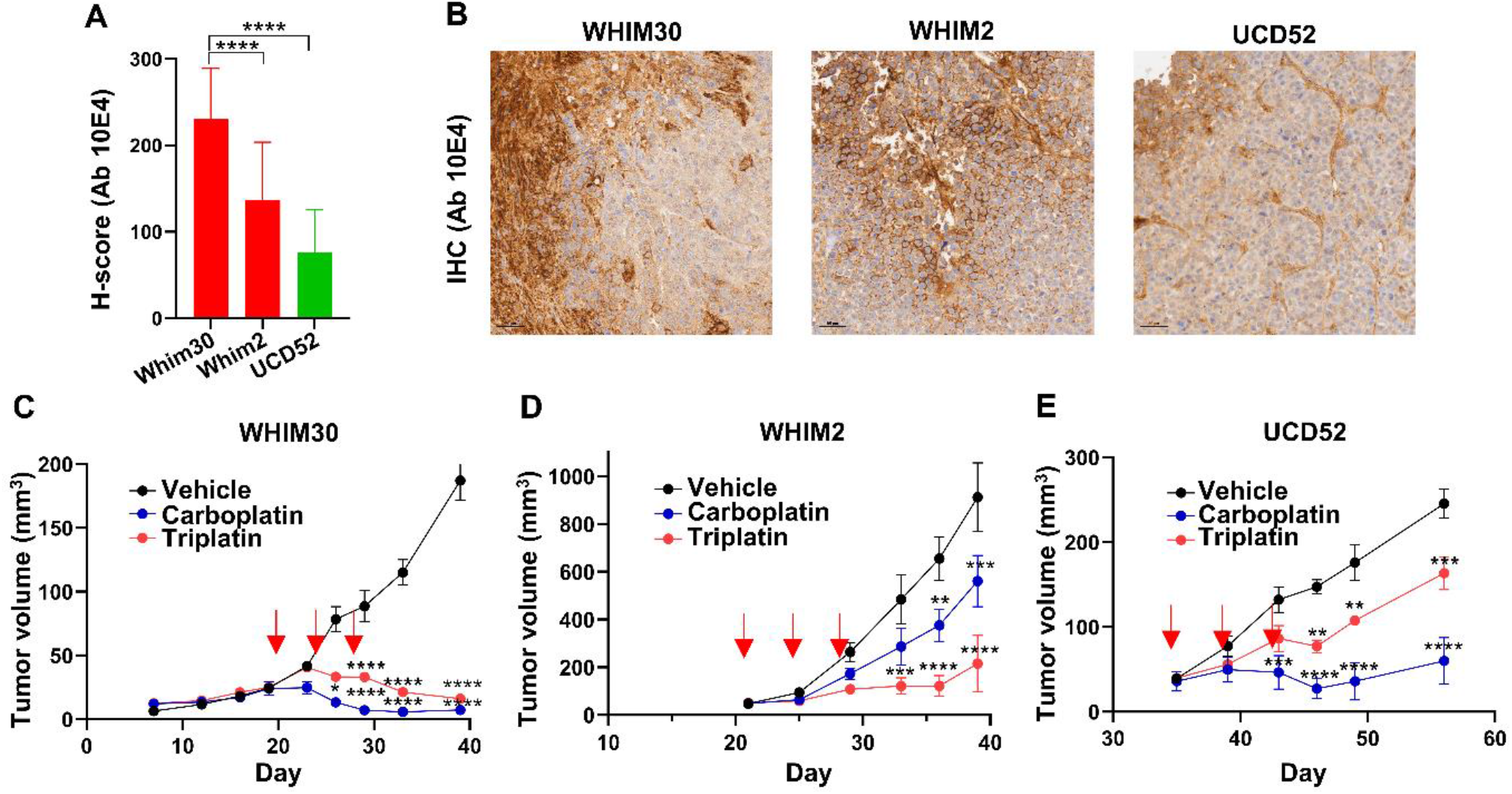
Triplatin efficacy against PDX TNBC primary tumor growth correlates with sGAG levels. **A** IHC H-scores of Ab 10E4 reactivity in TNBC PDX tumors. H-scores expressed as mean ± SD (bars). Statistical significance evaluated by one-way ANOVA (n=3). **B** Representative images of Ab 10E4 IHC staining. Scale bar represents 50 µm. (**C-E**) Analysis of primary tumor growth; 1×10^5^ cells injected into the T4 mammary gland of female NSG mice. Animals randomized and injected i.p. Q4Dx3 with Triplatin (0.3mg/kg), carboplatin (40 mg/kg), or vehicle (saline). Tumor growth measured by caliper and IVIS imaging. Statistical analysis: two-way ANOVA (n=5). *p<0.05, **p<0.01, ***p<0.001, ****p<0.0001

The effects of Triplatin and carboplatin on metastatic growth of WHIM30, WHIM2, and UCD52 PDXs were further evaluated using an intracardiac injection model. As previously described, WHIM30 cells formed metastases in the lung and liver. Similar to the primary tumor study, Triplatin and carboplatin decreased metastatic burden overall (Fig. 6A-B) and specifically, in the liver (Fig. 6C, 6E) and the lung (Fig. 6D). WHIM2 cells formed metastases in the brain, lung, liver, and ovaries with the highest burden in the lung and ovaries. Triplatin, but not carboplatin, decreased metastatic burden overall (Fig. 6A-B); specifically, in the ovaries (Fig. 6C, 5F) and the lungs (Fig. 6D). UCD52 formed metastases in the lung, liver, and brain with the highest burden in the liver and brain, which were overall more sensitive to carboplatin than Triplatin (Fig. 6A-B). In specific organs, carboplatin inhibited metastases to the liver more efficiently than Triplatin (Fig. 6C-D), and only carboplatin inhibited brain metastatic growth (Fig. 6D). Taken together, these results demonstrate that Triplatin may be more effective than carboplatin in treatment of TNBC with high sGAG levels..

**Figure 6.**
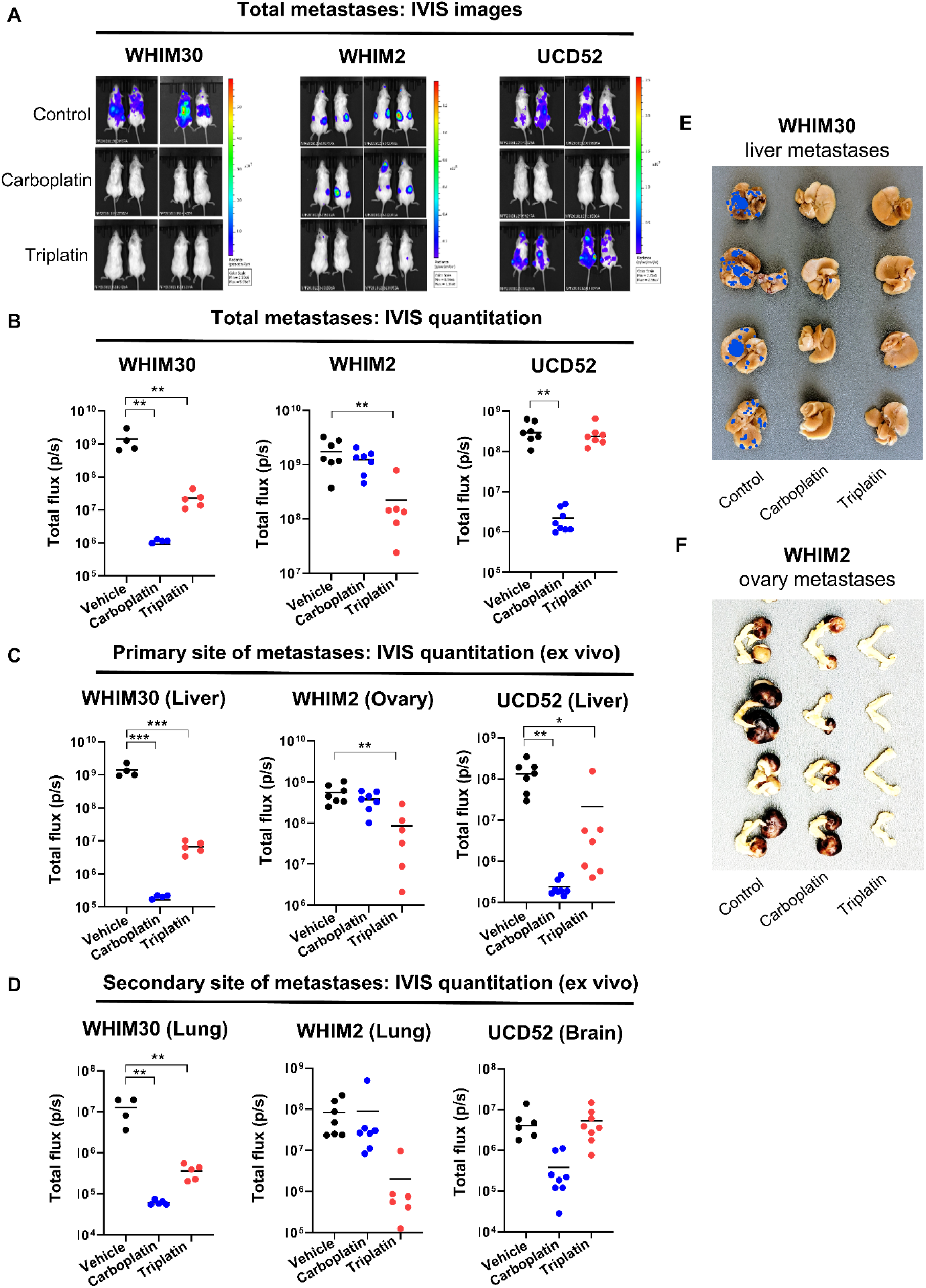
Efficacy of Triplatin against PDX TNBC metastatic growth correlates with sGAG levels. WHIM30, WHIM2, or UCD52 PDX cells (5×10^5^) were injected into left ventricle of female NSG mice. Animals were randomized and injected i.p. Q4Dx3 with Triplatin (0.3mg/kg), carboplatin (40 mg/kg), or vehicle (saline). **A** Total metastases were imaged *in vivo*. **B** Total metastases were quantified for each model. **C**-**D** Tumors and organs were harvested for *ex vivo* imaging and quantified. **E** Light photographs of WHIM30 liver metastases. Visible metastases highlighted using false coloration. **F** Light photographs of WHIM2 ovary metastases. Statistical significance was determined by one-way ANOVA, Tukey’s posttest. *p<0.05, **p<0.01, ***p<0.001, ****p<0.0001

## Discussion

Although numerous reports describe molecular characterizations of TNBC (including genetic mutations, copy number alterations and protein expression), quantitation of carbohydrate levels is generally omitted. However, a few pivotal studies have identified upregulation of HS and CS in BC compared to normal tissue, (Table 1). Gill et al, showed high expression of the HS 10E4 antibody epitope in clinical samples of BC and, in particular, TNBC (25). We applied this antibody to a TMA of TNBC PDX tumors and found distinct groups of low and high staining tumors. Tumors with H-scores >100 were predictive of Triplatin efficacy, even in models that were carboplatin-resistant.

The desirability of incorporating a platinum into treatment regimens stems from data suggesting that a high frequency of DNA damage repair defects renders TNBC particularly susceptible to DNA-modifying agents (8,40). Platinums exert their effect by creating DNA-drug lesions that inhibit DNA replication and induce cell death. The synergy of two different mechanisms that potentially induce DNA damage (platinums causing DNA damage and “BRCAness” with ineffective DNA damage repair), known as synthetic lethality, is the rationale for the potential benefit of platinums in this setting. Triplatin was originally developed to generate DNA lesions that are distinct from other platinums and less recognizable by DNA repair proteins. Indeed, the 1,4 DNA interstrand crosslinks produced by Triplatin, unlike the 1,2 intrastrand crosslinks produced by cisplatin/carboplatin, evade binding of high mobility group proteins and repair by the nucleotide excision repair machinery (41–43). The results shown here are consistent, in that Triplatin showed similar activity in high sGAG-expressing PDX models, WHIM30 (mtBRCA) and WHIM2 (wtBRCA), regardless of BRCA status, whereas carboplatin was markedly less effective in the WHIM2 model with wtBRCA (36,37). Together, these results suggest that the efficacy of Triplatin may be independent of BRCA status.

The majority of TNBC patients in the USA present with local or locally advanced (stage I-III) disease at the time of diagnosis. Patients who achieve pCR, defined as elimination of invasive cancer from the breast and lymph nodes, have excellent chances of recurrence-free survival. However, those with residual disease (RD) have a higher risk of relapse and poorer prognosis than other BC subtypes. Paradoxically, a subset of TNBC appears to be particularly chemosensitive, with 30-40% achieving pCR after NACT using standard anthracycline and taxane-based combinations. Four recent randomized trials, GeparSixto, CALGB 40603, Brightness and I-SPY 2, have investigated the addition of carboplatin to standard NACT regimens. On average, the pCR rate increased to 50-55% when carboplatin was included (4,30,44–46). These results were confirmed in the ongoing Keynote-522 trial and with the addition of pembrolizumab, the number of patients achieving pCR further increased to ~65% (1). Together, the results support adding platinum to NACT regimens in treatment of locally advanced TNBC, which may again be optimized in the context of BRCA status and/or homologous repair deficiencies (HRD).

The major advantage of NACT is the ability to immediately identify patients with chemotherapy-resistant disease who may benefit from second adjuvant therapy following surgery (47). Recent approval of capecitabine, an oral antimetabolite of 5-Flurouracil, resulted from the CREATE-X trial showing a significant improvement in the rate of recurrence-free survival at 5 years (69.8% in the capecitabine group vs 56.1% in the control group) (48). The ongoing EA1131 trial will challenge this success by comparing capecitabine directly with 6 cycles of carboplatin (4). Future studies profiling residual disease for markers of selective targeting agents, such as Triplatin, may further benefit patients with chemotherapy-resistant disease.

Patients who developed distant metastases during their disease survive less than a year, underscoring the importance of developing more effective treatment of patients living with metastatic TNBC. Even with the addition of recent treatment options for metastatic TNBC, such as immunotherapy (nab-paclitaxel and atezolizumab) for PD-L1-positive tumors and carboplatin/PARP-inhibitors for BRCAness-positive tumors, the survival benefit is measured in months (6,8,49). For this reason, the inclusion of patients in ongoing trials evaluating new targeted agents and predictive biomarkers is typically encouraged. The results presented here find that Triplatin is effective at prolonging survival of mice with metastatic TNBC and inhibiting growth of TNBC cells at metastatic sites in TNBC with high levels of sGAGs in multiple models.

In light of our discovery that Triplatin cellular and tumor uptake is sGAG-dependent, a reevaluation of the clinical potential of Triplatin in this context is warranted. The primary dose-limiting toxicities (DLT) of Triplatin were neutropenia and diarrhea. Using the recommended Phase II dosing schedule, 1.1 mg/m^2^ on d1 of a 28-day cycle, Triplatin had a manageable toxicity profile. Phase I trials showed objective responses in advanced metastatic BC; in combination with 5-FU (1 partial response, 3 stable disease/14) (50). Phase II clinical trials in cisplatin-relapsed ovarian cancer gave partial responses (5/28 patients), (43) and in several cases durable responses > 3 years (personal communication).

In summary, these results show activity of Triplatin in TNBC tumors with high sGAG levels, regardless of BRCA status and carboplatin-resistance. Although the added toxicities of incorporating platinum into any regimens should be considered, the benefits may be substantial, especially in early stage cases showing resistance, residual disease after neoadjuvant therapy, and in the metastatic setting. Elevation of pCR rate is expected to significantly improve patient survival.

### Respective Contributions of Authors (in alphabetical order of surname)

Original idea and project development-NPF, JDH, SJK, JEK, EJP, KT; acquisition of data including provision of cell lines - MAZ, PG, BH, JCH, JDH, EK, JEK, SJK, VK, EJP, MP, MS, THT; discussion, analysis and interpretation of data-MD, JCH, JDH, MI, JEK, SJK, LL, EJP, KT; writing-NPF, JDH, JEK, SJK, EJP. All authors have reviewed and approved this manuscript.

## Acknowledgements

We thank Dr. Jeff Esko for the CHO-F17 mutant cell line and Dr. Harry Bear for the use of 4T1 tumor samples for IHC. The lentivirus stocks have been generated by Gene Editing, Transduction and Nanotechnology Core, Skin Biology & Disease Resource-Based Center, Northwestern University *supported in part by the NIH-NCI Cancer Center Support Grant (P30 CA060553)*. Services in support of the research project were provided by the Virginia Commonwealth University Massey Cancer Center Flow Cytometry Core, Cancer Mouse Models Core Laboratory, and Microscopy Facility supported, in part, with funding to the Massey Cancer Center from NIH-NCI Cancer Center Support Grant P30 CA016059. This work was supported by the Massey Cancer Center Molecules to Medicine Initiative, Commonwealth Health Research Board (CHRB 236-05-17), Commonwealth Research Commercialization Fund (MF17-028-LS) and Value and Efficiency Teaching and Research Initiative (VETAR, School of Medicine, VCU). National Institutes of Health Grant R01CA160688 to KT. National Cancer Institute (NCI) Grants P30CA016056 and U24CA232979, involving the use of Roswell Park Comprehensive Cancer Center’s Shared Resources for KT and EK.

## Conflict of Interest Statement

The authors declare no potential conflicts of interest.

## Notes

### Competing Interest Statement

The authors have declared no competing interest.

