## Supplemental Information for "Exploitation of sulfated glycosaminoglycan status for precision medicine of platinums in triple-negative breast cancer"

### Supplementary Methods

#### *IHC*

All tissues were fixed in 10% neutral buffered formalin for at least 5 days before paraffin embedding and sectioning. A TMA was obtained from Champions Oncology containing breast cancer PDX samples. IHC staining for total heparan sulfate (Amsbio, heparan sulfate neo epitope clone F69-3G10, 1:100 dilution), was performed in the VCU Cancer Mouse Models Core with the Leica Bond RX autostainer. All antibodies were incubated for 30 min at 25ºC. Positive slides for total heparan sulfate staining (3G10) were incubated with 50 mU Heparinase III (Sigma-aldrich) n Tris buffer (20 mM Tris-HCL, pH 7.5 containing 0.1 mg/ml bovine serum albumin and 4 mM $CaCl_2$) for 45 min at 37ºC, 3 times, before primary antibody incubation. Digestion with heparinase III cleaves the heparan sulfate chain that exposes a neo-epitope for 3G10. For negative total HS staining (3G10) slides were incubation with only Tris Buffer for 45 min at 37ºC, 3 times, before primary antibody incubation. Bond polymer refine detection kit was used for DAB chromogen and hematoxylin counterstain (Leica). The tissue slides were visualized with a Vectra® Polaris™ Automated Quantitative Pathology Imaging System (Akoya Biosciences) using whole-slide scanning at 20x. Phenochart Whole Slide Contextual Viewer software was used to visualize the scans and inFrom software (Akoya Biosciences) was used to quantify the staining. The wt CHOK1 and psA-745 were used to titrate the 3G10 antibody and validate staining.

#### *Syngeneic orthotopic mouse model*

Female BALB/c mice were orthotopically injected with 4T1-*luc2* cells ($1x10^4$ cells suspended in 20µL 1:9 mix of PBS:Cultrex BME Type 3 [14 mg/ml, Biotechne]) into 2nd mammary fat pad on day 0. Primary tumor was surgically removed on day 8 and mice were then randomized into treatment groups. The mice were treated with Triplatin (0.3 mg/kg) or saline control on days 10, 14, and 18 by i.p. injections. For endpoint: on day 21, mice were injected with D-Luciferin (150 mg/kg diluted in PBS) allowing for live *in vivo* imaging using the IVIS Spectrum and Living Image® software (PerkinElmer). After imaging mice were euthanized. Lungs were harvested for *ex vivo* bioluminescent imaging immediately after sacrifice and analyzed using the photon emission with Living Image Software (Xenogen, PerkinElmer). n = 6 mice, each group

#### *Syngeneic Metastasis mouse model*

BALB/c female 6-8-week-old mice were injected into the left cardiac ventricle with $4x10^4$ murine mammary cancer 4T1-*luc* cells (in 100 µL PBS) on day 0. Mice were randomized on day 1 based on total flux values and body weight into 3 groups (Triplatin, carboplatin, control). Then the mice were treated by i.p. with Triplatin (0.3 mg/kg), carboplatin (40 mg/kg), or saline control on days 1, 5, and 9. Tumor burden was quantified by bioluminescence (radiance/sec) emitted from the cells after a 200 µL subcutaneous injection of D-luciferin (150 mg/mL diluted in PBS) allowing for live *in vivo* imaging using IVIS Spectrum and Living Image® software (PerkinElmer). On day 13, mice were euthanized and organs (kidneys, heart, lung, ovaries, liver, brain, and skeleton) were harvested and immersed in luciferin/PBS (3 mg/ml) for 5 min before *ex vivo* imaging with the IVIS Spectrum. n = 10 mice, each group

### Supplementary Figures

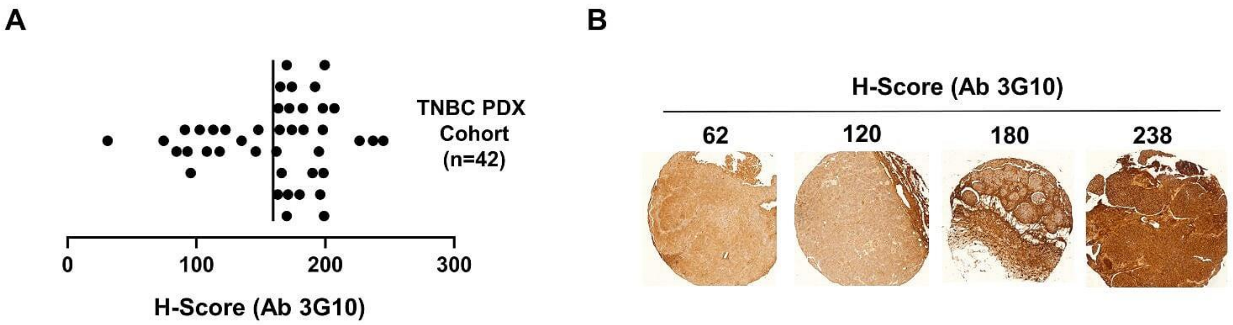


**Supplementary Figure 1. 3G10 PDX TMA Staining A.** IHC H-scores of TNBC PDX samples on TMA (n=42) stained with Ab 3G10. Each data point represents the mean H-score for duplicate samples. **B.** Representative images of Ab 3G10 staining of TNBC PDX TMA samples.

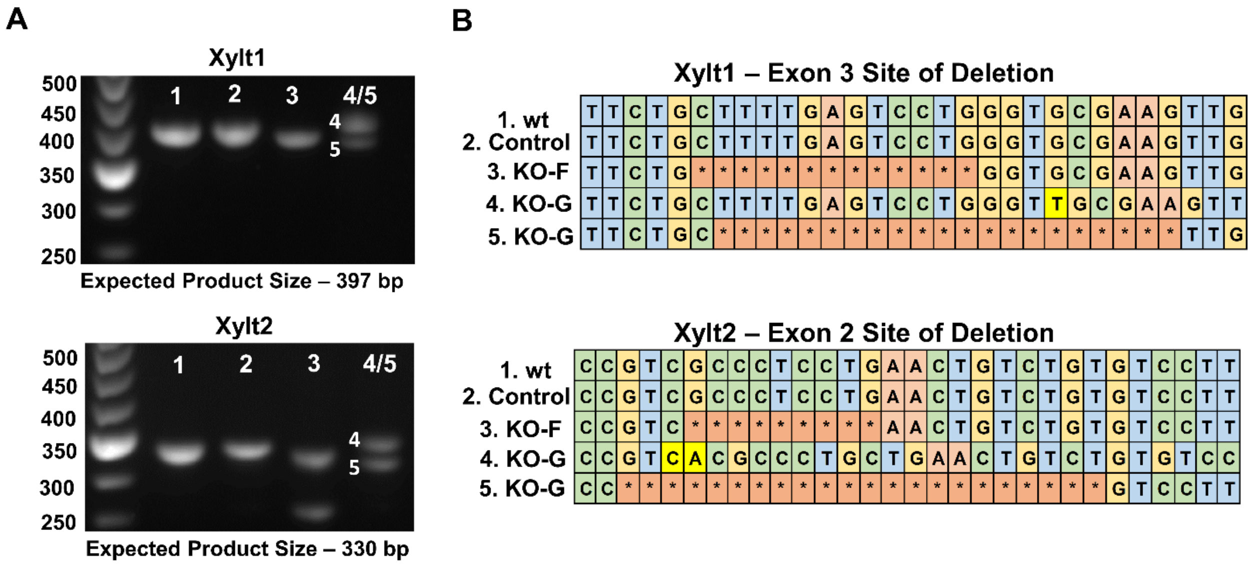


**Supplementary Figure 2. MDA-MB-231 XYLT1/2 KO confirmation. A.** DNA Gels showing generated PCR products for MDA-MB-231 wt (1), CRISPR Control (2), XYLT1/2 KO-F (3) , XYLT1/2 KO-G (4-5). **B.** Sequences of generated PCR products from MDA-MB-231 XYLT1/2 KO. Red squares with astericks show location of deletions within product. Bases in yellow are insertions.

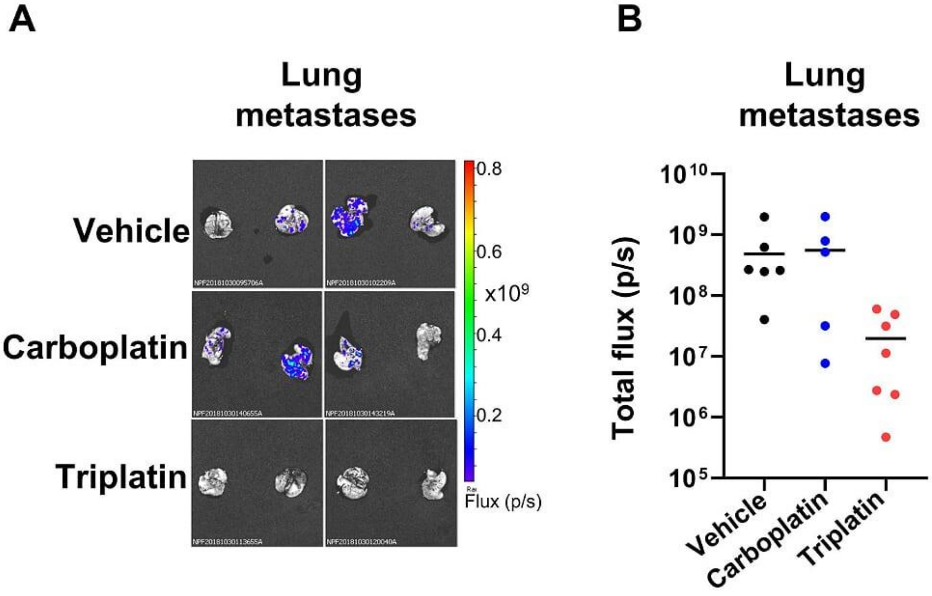


**Supplementary Figure 3. MDA-MB-231-luc lung metastases photos and quantitation** MDA-MB-231BR-luc cells ($1x10^4$) were injected into left ventricle of female NSG mice (day 0). Animals were randomized on day 10. Drug treatment [Triplatin (0.3mg/kg) or control (saline)] were given i.p. on days 10, 14, and 18. Animals were euthanized on day 21. Tumors and organs were harvested for *ex vivo* imaging and analysis. **A.** Images of lung metastases *in vivo* on day 21 (total flux). **B.** Quantification of lung metastases *ex vivo* on day 21 (total flux). Statistical analysis: One-way ANOVA; no statistical differences

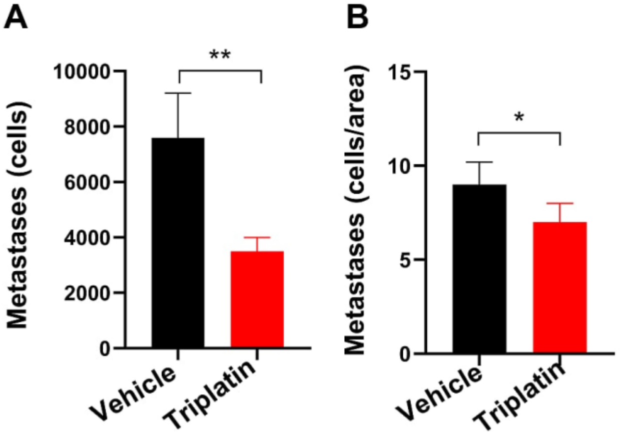


**Supplementary Figure 4. MDA-MB-231-BR intracardiac model – endpoint; liver metastasis quantitation** Quantification of liver metastases *ex vivo* on day 21 using HLA staining. Cell Number (**A)** and Cells/area (**B)** were determined using Image J software. Statistical analysis: Student unpaired t-test *p<0.05, **p<0.01

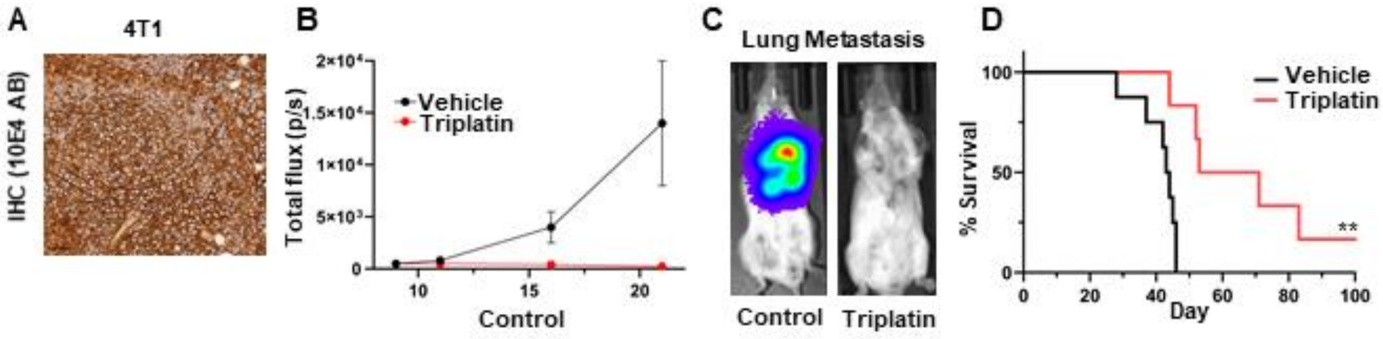


**Supplementary Figure 5. 4T1 mastectomy model – lung metastasis** Analysis of lung metastasis and survival after primary resection; 4T1-luc cells (1x$10^5$) were injected into mammary gland of female BALB/c mice (day 0). Primary resection was performed on day 8 and animals were randomized. Drug treatment [Triplatin (0.3mg/kg) or control (saline)] was given i.p. on days 10, 14, and 18. n=6, each group**. A.** Representative images of Ab 10E4 IHC staining. Scale bar represents 50 µm. **B.** Quantification of lung metastasis *in vivo* (total flux) **C.** Images of lung metastases *in vivo* (IVIS) on day 21. **D.** Survival analysis. Statistical analysis: Student unpaired t-test *p<0.05, **p<0.01, ***p<0.001, ****p<0.0001

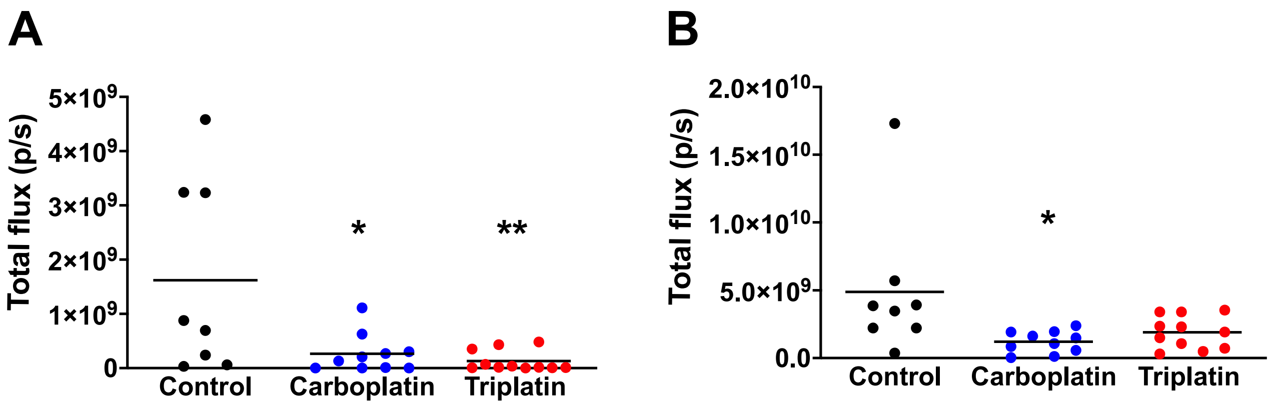


**Supplementary Figure 6. 4T1 intracardiac model – endpoint;** Analysis of lung and bone metastasis. 4T1-luc cells (4x$10^4$) were injected into the left ventricle of female BALB/c mice (day 0). Animals randomized on day 1. Drug treatment [Triplatin (0.3 mg/kg), carboplatin (40 mg/kg) or control (saline)] given i.p. on days 1, 5, and 9. n=10, each group**.** Animals sacrificed on day 21. Tumors and organs harvested for ex vivo imaging and analysis. Quantification of **A.** Lung and **B.** Bone metastases *ex vivo* on day 13 (total flux). Statistical analysis: One-way ANOVA; Tukey's posttest *p<0.05, **p<0.01
